# Comparative Analysis of rRNA removal methods for RNA-seq Differential Expression in Halophilic Archaea

**DOI:** 10.1101/2022.04.15.488396

**Authors:** Mar Martinez-Pastor, Saaz Sakrikar, Deyra N. Rodriguez, Amy K. Schmid

**Affiliations:** Biology Department, Duke University, Durham, NC, 27708, USA; University Program in Genetics and Genomics, Duke University, Durham, NC, 27708, USA; New England Biolabs, Ipswich, MA, 01938, USA

## Abstract

Despite intense recent research interest in archaea, the scientific community has experienced a bottleneck in the study of genome-scale gene expression experiments by RNA-seq due to the lack of commercial and specifically designed rRNA depletion kits. The high ratio rRNA:mRNA (80-90%: ∼10%) in prokaryotes hampers global transcriptomic analysis. Insufficient ribodepletion results in low sequence coverage of mRNA and therefore requires a substantially higher number of replicate samples and/or sequencing reads to achieve statistically reliable conclusions regarding the significance of differential gene expression between case and control samples. Here we show that after the discontinuation of the previous version of RiboZero (Illumina) that was useful to partially deplete rRNA from halophilic archaea, archaeal transcriptomics studies have experienced a standstill. To overcome this limitation, here we analyze the efficiency for four different hybridization-based kits from three different commercial suppliers, each with two sets of sequence-specific probes to remove rRNA from four different species of halophilic archaea. We conclude that the key for transcriptomic success with the currently available tools is the probe-specificity for the rRNA sequence hybridization. With this paper we provide insights to the archaeal community for selecting certain reagents and strategies over others depending on the archaeal species of interest. These methods yield improved RNA-seq sensitivity and enhanced detection of low abundance transcripts.

## INTRODUCTION

The expression of the genomic information of an organism depends on the cell status and environmental factors that determine the phenotype. The genes across the genome that are being transcribed collectively define the transcriptome, and the compendium of methods that enable the study of the expression of large number of genes simultaneously is known as transcriptomics. RNA sequencing (RNA-seq) has emerged as a widely used approach to transcriptome profiling with high throughput, sensitivity, dynamic range, and relatively low cost compared to former methods such as microarrays. The first successful RNA-seq experiments were performed using eukaryotic model organisms [1-4]; however, using this tool for understudied models such as archaea has been challenging despite their biological and evolutionary importance.

Archaea are prokaryotic microorganisms that were defined as the third branch of life in the late 70’s, when Carl Woese and colleagues found substantial 16S differences that warranted classifying the Archaea as a distinct group separate from the Bacteria and Eukarya [5]. Recent phylogenetic evidence is more consistent with a two-domain tree, with Eukarya stemming from Archaea [6]. Although archaea are typically known for their survival in extreme environments, archaeal species are now known to be diverse and abundant, colonizing a vast array of habitats (from oceans to human skin to extreme environments [7,8]). Therefore, differential expression analysis using transcriptomics in archaea is an important step for a better understanding of responses to diverse environments [9,10]. Such studies advance knowledge of the unique molecular biology of archaea, which combine the molecular characteristics of both bacteria and eukaryotes, such as transcriptional regulation [11].

Despite previous progress on differential expression by RNA-seq in archaea [12], this method has recently become unavailable. Previously, archaeal transcriptomics studies successfully depleted rRNA using commercially available reagent kits for rRNA removal in bacteria [13-17]. However, these kits were discontinued in 2018. Ribosomal RNA (rRNA) in archaeal transcriptomes can reach more than 90% of the total cellular RNA. As we report in the current work, ribodepletion is a key step for reliable RNA-seq results because high rRNA sequencing reads can preclude the detection of messenger RNA (mRNA). rRNA removal enables higher sequencing depth of mRNA, leading to better detection of transcripts. This is critical for analyzing differential expression, particularly when detecting non-coding or lowly expressed RNAs [18]. Previous studies have suggested a minimum sequencing depth of two [19] to ten [20] million reads per sample for obtaining reproducible results for differential expression, while the ENCODE consortium [21] mandates 30 million reads (albeit for much larger human genomes). Such sequencing depth enables sound statistical comparisons of differential expression on a per-gene basis: at least 5 reads per gene are typically needed to detect the significance of change in expression for a given gene [18]. Some archaeal studies have reported RNA-seq without rRNA removal, but these were conducted for different purposes that are possible without rRNA removal (e.g. transcription start site mapping [22], small RNA detection [12], etc). Removing rRNA also substantially reduces the cost of RNA-seq, enabling extensive sample multiplexing in a single sequencing run, especially for relatively small archaeal genomes (∼2-8 Mbp).

In this work, we have studied four species of halophilic archaea that have been widely used as model organisms in the archaeal research community: *Halobacteium salinarum* (HBT) and *Haloarcula hispanica* (HAH) of the family Halobacteriaceae require salt concentrations close to saturation, whereas *Haloferax volcanii* (HVO) and *Haloferax mediterranei* (HFX) of the family Haloferacales colonize lower salinity environments. These four species are highly tractable models for extremophilic microorganisms given their relatively fast generation time (2-6 hours in rich medium), facile genetic tools [23-25], and highly curated genomic annotations and databases [26-29]. Establishing a set of tools and best practices for transcriptomics methods would therefore greatly facilitate advances in this field.

Archaeal RNA, like that of bacteria, lacks a 3’ polyA tail, so rRNA cannot be removed by polyT tagging. Here we test two methodologies for rRNA depletion in archaea using: (a) biotinylated probes; and (b) enzymatic digestion. We use probes that come packaged with commercial kits as well as sequence-specific probes customized for particular species of interest. The first approach (biotinylated probes/streptavidin beads) consists of a physical removal of rRNA by hybridizing with a pool of biotinylated oligo probes. These probes are then be captured and removed from the RNA sample using streptavidin-coated magnetic beads. In contrast, the enzymatic removal of rRNA consists of generating DNA-rRNA hybrids by incubating specifically designed DNA probes complementary to rRNA. Hybrids are then treated with RNaseH that catalyzes the cleavage of RNA when it is bound to a DNA substrate.

Here we report that the two methods are equally successful for removing rRNA across the four species of halophilic archaea growing in diverse media. Both methods can be used successfully with probe sequences custom-designed for one species or with a broad probe pool designed to target multiple species simultaneously. We show that bacterial rRNA probes are sufficiently divergent in sequence to preclude the use of recently developed custom and commercial bacterial rRNA probe sets in archaea [30]. These methods are robust to varying culturing conditions (rich and defined media). This analysis has achieved the goal of identifying an efficient and broadly useful strategy for depleting undesirable archaeal rRNA prior to sequencing for successful transcriptomics.

## MATERIAL AND METHODS

### Media, Strains, and Growth Conditions

All used strains, media and conditions for this study are summarized in the following **Tables 1** and **2**.

**Table 1:** Strains used in this study.

| Name | Species abbreviation | Genotype | Reference genome |
| --- | --- | --- | --- |
| MDK407 [24] | HBT | Δ*ura3* | GCF_000006805.1_ASM680v1 |
| DS2 [31] | HVO | Δ*pyrE* | GCF_000025685.1_ASM2568v1 |
| ATCC33500 [25] | HFX | Δ*pyrE* | GCF_000306765.2_ASM30676v2 |
| DF60 [25] | HAH | Δ*pyrF* | GCF_000223905.1_ASM22390v1 |

**Table 2:**
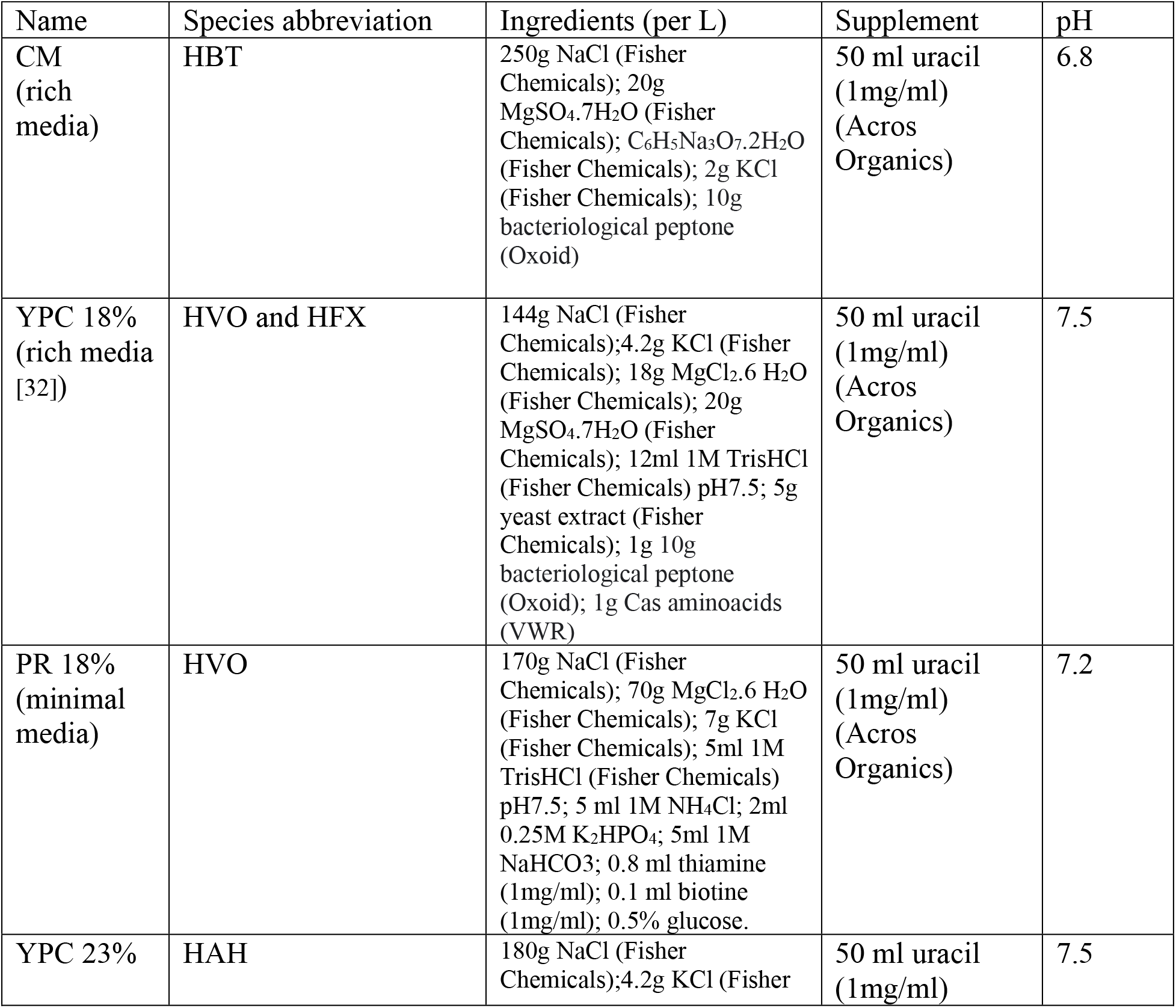
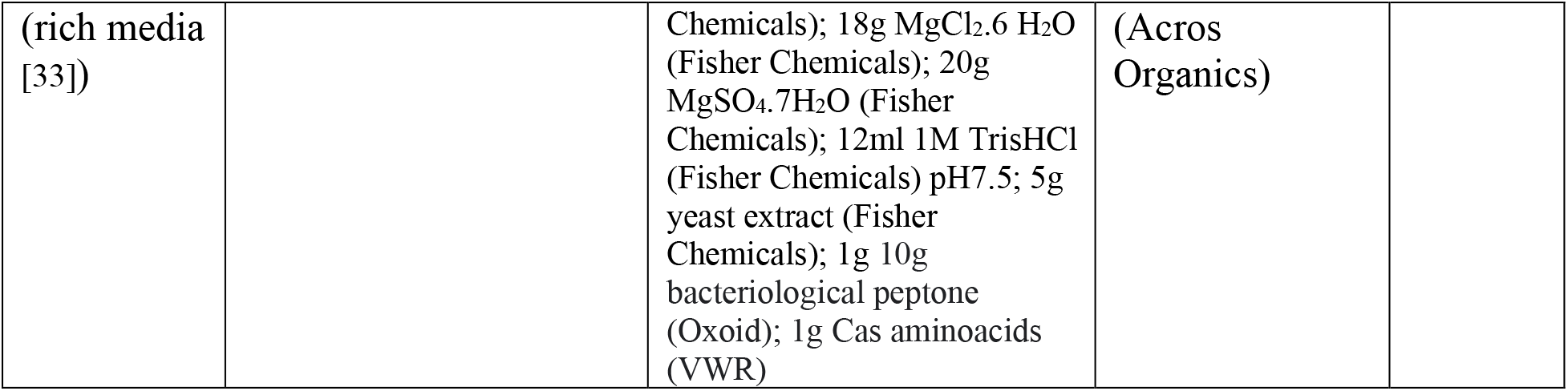
All media recipes used for test organisms in this study

For routine culturing in these media, each species was freshly streaked from frozen stock. Single colonies from no more than two-week-old plates were inoculated in triplicate in 3 ml of rich, minimal, or defined liquid media (**Table 2**) and grown aerobically until saturation (stationary phase) at 42°C with continuous shaking at 225 rpm. From each saturated pre-culture, 50ml cultures were initiated by diluting the pre-culture to OD_660_=0.1 in 150 ml Pyrex flasks, and 3ml of each were harvested in mid exponential phase OD_660_=0.4-0.8 (doubling times and incubation times included in **Table 3**), by centrifugation in a tabletop centrifuge (5424, Eppendorf) at 21,130 x g for 3min. Supernatant was discarded and pellets were immediately snap-frozen in liquid N_2_ and stored no longer than 3 weeks at -80°C until RNA extraction.

**Table 3:** Doubling time and incubation time for different species in different media.

| Species | media | doubling time (h) | days until stationary phase |
| --- | --- | --- | --- |
| HBT | CM | 6 | 3 |
| HVO | YPC18% | 3 | 2.5 (36h) |
| HVO | PR18% | 12 | 3 |
| HFX | YPC18% | 2.5 | 2 |
| HAH | YPC23% | 6 | 3 |

### RNA-seq experimental protocol

Total RNA was extracted from pellets using Absolutely RNA Miniprep kit (Agilent Technologies, Santa Clara, CA) according to manufacturer’s instructions. The obtained RNA concentration and integrity was quantified by Nanodrop One (Thermo Scientific, Grand Island, NY) and RNA electropherograms, Bioanalyzer 2100 Instrument with the RNA 6000 Nano kit (Agilent Technologies, Santa Clara, CA), respectively. RNA was checked for DNA contamination by PCR using 200-300ng of input RNA and primers given in **Table 4** for 30-35 cycles. Extracted RNA was high quality in all samples, with Bioanalyzer RNA integrity number (RIN) greater than 8.

**Table 4:**
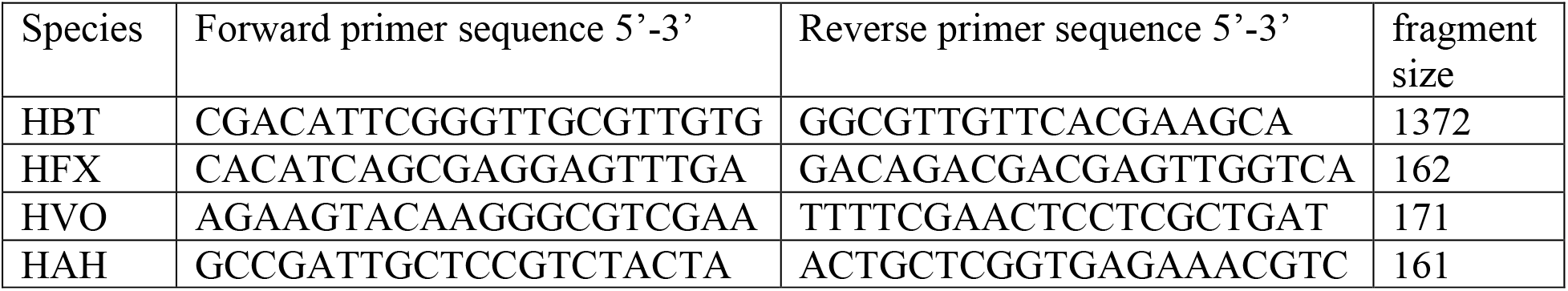
Primers used to check for genomic contamination

Ribosomal RNA was removed using the following reagent kits and methodologies, abbreviated throughout the text and figures as indicated below:

1. Biotinylated probes with strepdavidin bead pull-down:
  a. Discontinued Ribo-Zero rRNA Removal Kit (Bacteria). Abbr: RZ
  b. siTools HVO RiboPOOL™ with probes specific for HVO. Abbr: rP-HVO
  c. siTools Pan-Archaea riboPOOL™ (probes included). Abbr: rP-PA
2. RNAse H and enzymatic depletion-based protocols with magnetic bead pull-down:
  a. Ribo-Zero Plus Kit (probes included). Abbr: RZ+
  b. NEBNext Bacteria rRNA depletion Kit (New England Biolabs) with probes designed for bacteria (included in kit from NEB). Abbr: NEB-B
  c. NEBNext Depletion Core Reagent Set with customized sequence-specific probes for HVO (Table S3). These probes were designed using the NEB web tool (https://depletion-design.neb.com/) and ordered from IDT techonologies (idtdna.com). Abbr: NEB-HVO

RNA input to each depletion kit was 300-500 ng. Ribodepletion was performed according to the manufacturer’s manuals using default or custom-designed probes as well as modifying time of enzymatic incubation with RNaseH. These details and ordering information are specified in **Table S1**.

Library preparation from 1-10ng rRNA-depleted RNA was performed using NEBNext UltraII Directional RNA Library Preparation Kit (Illumina, #E7760) following the vendor protocols and complementing cleaning steps with NEBNext Sample Purification Beads (#E7767). An extra-cleaning step using the same type of beads was carried out when samples showed contamination with adaptor dimers. The obtained library quality and concentration was assessed by monitoring the distribution of the fragment sizes with a Bioanalyzer 2100 instrument using RNAnano reagent kit (Agilent Technologies, Santa Clara, CA). This size and quantity information was used for pooling the libraries in equimolar concentrations to normalize each library. Libraries were subjected to HiSeq2500, HiSeq4000, or NovaSeq6000 by the Sequence and Genomics Technologies Facility at Duke University. Additional experimental metadata, results, and details are given in **Table S1**.

## Data Analysis

### Publications on archaeal RNA-seq per year

Data regarding the number of publications yearly available from National Center for Biotechnology Information (NCBI) PubMed database (https://pubmed.ncbi.nlm.nih.gov/) was searched with the phrases “archaea”, “RNA-Seq”, and “archaea RNA-Seq” Database hits were downloaded from the NCBI PubMed database on November 1, 2021. The publication of Carl Woese’s seminal paper regarding the classification of Archaea in 1977 [5] was used as the starting date. The downloaded data is in **Table S2**. The code used to generate **Fig. 1** is in https://github.com/amyschmid/rRNA_analysis.

**Fig 1.**
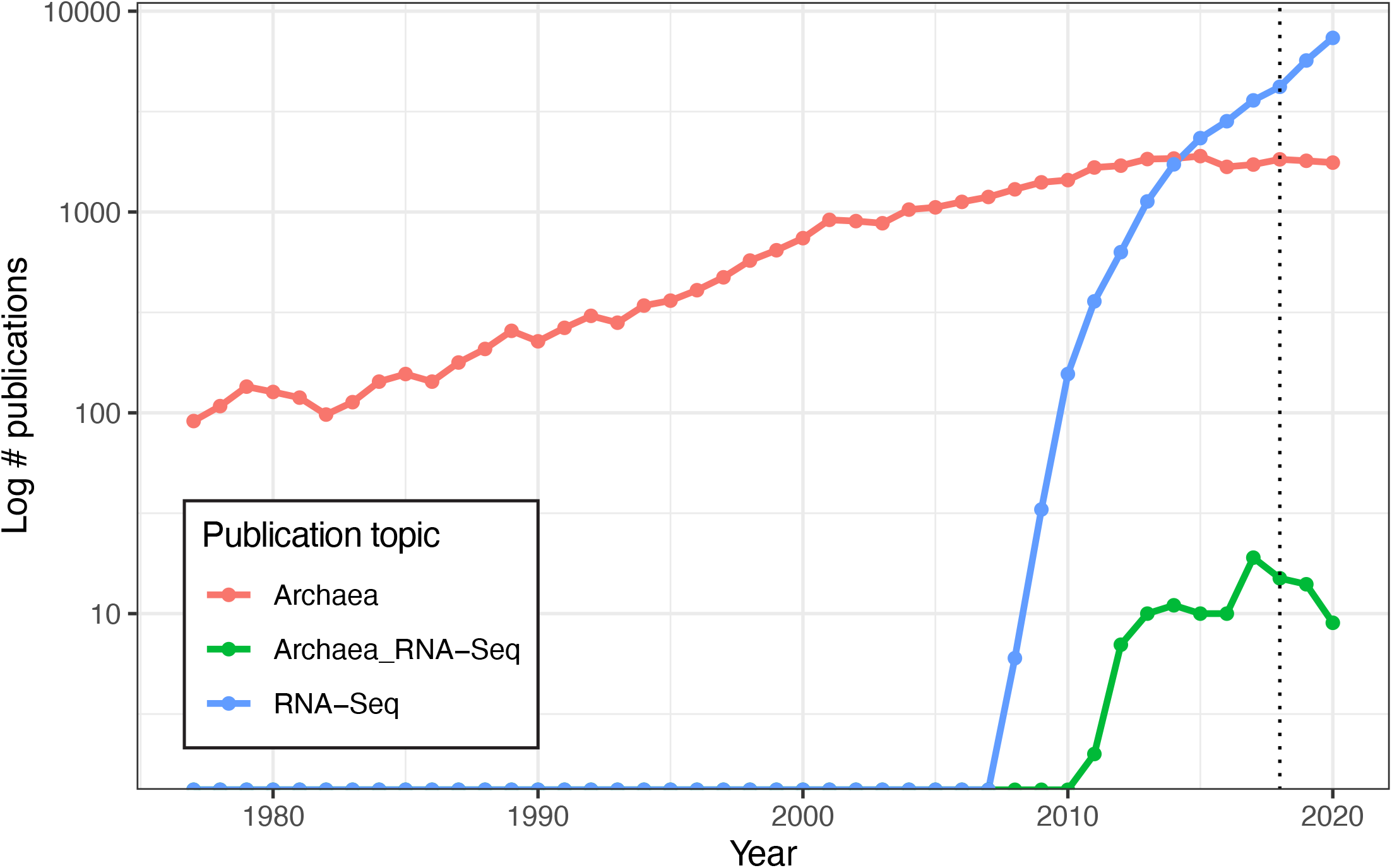
Slowdown in Archaeal RNA-Seq publications in recent years. Lines depicting number of publications per year detected in the NCBI PubMed databased searched with the terms “Archaea” (red), “RNA-Seq” (blue), and “Archaea RNA-Seq” (green), plotted on log-scale y-axis. Dotted line at 2018 marks discontinuation of the Illumina RiboZero kit.

### RNA-seq data processing

FASTQ files generated by sequencing were downloaded and processed as described previously[34]. Files were quality-checked using FastQC, adapter sequences were trimmed using TrimGalore! with cutadapt (FastQC and TrimGalore! downloaded from http://www.bioinformatics.babraham.ac.uk/projects/). Trimmed files were aligned to the reference genomes of the four species of interest (Table 1) using Bowtie2 [35]. Resultant SAM files were converted into a compact BAM file using SAMtools [36] to generate, sort, and index reads. BAM files were used as the input for HTSeq-count [37] to generate a count file, assigning a numeric raw count of reads to each gene. Details regarding the full workflow are included in reference [34]. To determine rRNA percentage remaining following depletion, the counts corresponding to each of 16S, 23S, and 5S rRNA genes was divided by the total number of raw counts mapping to all genes. The ratio was multiplied by 100 to yield a percentage. These genes are listed in **Table 4**.

**Table 4.**
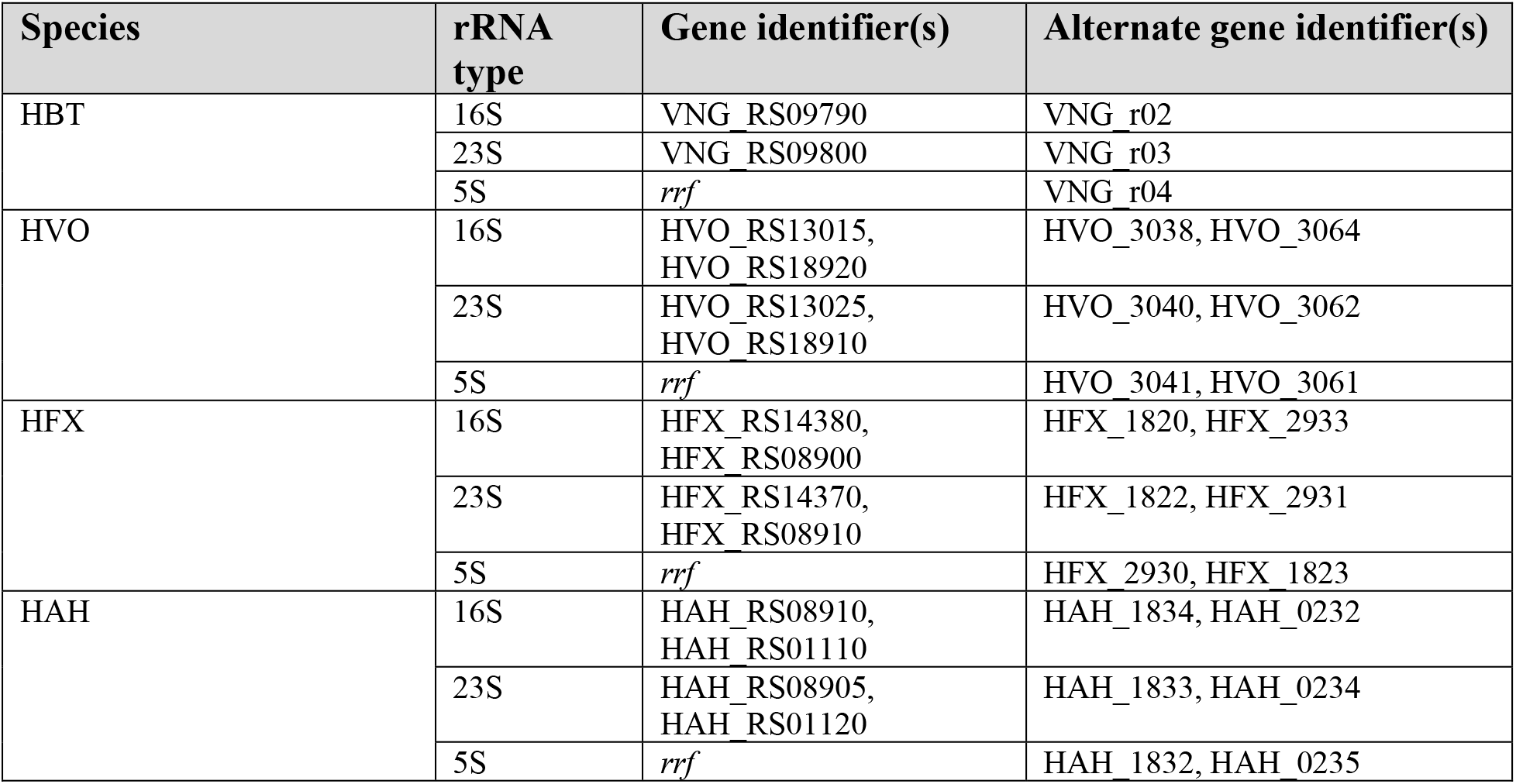
rRNA-coding gene identifiers for each species of interest.

The results, expressing all rRNA, 16S rRNA, 23S rRNA, and 5S rRNA as a percentage of total reads, are listed in **Table S1**. The code used to generate **figures** is in https://github.com/amyschmid/rRNA_analysis, and the input to the code is also given in **Table S1** under the appropriate tabs.

### Probe specificity analysis

Sequences of probes custom-designed for HVO rRNA removal using the NEB website (https://depletion-design.neb.com/) were compared to HBT strain NRC-1 genome sequence using NCBI BlastN search with default parameters (NCBI taxonomy ID: 64091; NCBI access date May 4, 2021). The resultant sequence identity (expressed as a percentage) was noted for each of the 117 sequences. These data were classified into 4 categories: 100% identity, 90-99% identity, <80% identity, and no significant similarity. The probe sequences, BLAST results, and identity percentages are listed in **Table S3**. The code used to generate the corresponding figure is in https://github.com/amyschmid/rRNA_analysis and the specific inputs to generate this figure are in the appropriate tabs within **Table S3**.

### Count correlations

RNA-seq read counts corresponding to all genes outside of rRNA genes for different rRNA removal methods and replicates in HBT and HVO were calculated as described above. Each gene’s count was expressed as a percentage of total counts, and the arithmetic average of all replicates using a particular method was calculated. These average values for each gene for a given removal method were then noted in **Table S4**. The code used to generate the corresponding figure is in https://github.com/amyschmid/rRNA_analysis and the specific inputs to generate this figure are in the appropriate tabs within **Table S4**.

### Power analysis

RNA-Seq data generated from a pilot run for a published project [34] from the Schmid lab was inputted into the power optimization tool Scotty (scotty.genetics.utah.edu) [38]. This was used to assess power for differential expression experiments involving up to 6 biological replicates with between 1 and 15 million reads mapping to genes for each replicate, so that at least 75% of 2-fold differentially expressed genes could be detected at *p*<0.01.

## RESULTS

### Discontinuation of the Illumina RiboZero kit is associated with a decline in published archaeal RNA-Seq studies

RNA-Seq of archaeal species belonging to diverse clades has previously been facilitated by rRNA depletion using the bacterial Ribo-Zero kit from Illumina [13-17] (Methods). However, the kit was discontinued in 2018. To determine the impact of this discontinuation, we conducted a comprehensive literature search on the PubMed database for articles reporting on archaea (1977-present) and on RNA-seq in archaea (2010-present). The discontinuation of the Ribo-Zero kit appears to correlate with a plateau and decline of papers published on the topic of RNA-seq in archaea, even as the number of publications on archaea in general and on RNA-Seq in other domains of life has grown (**Fig. 1**). Within our lab, we had successfully used this kit on two model halophile species, HBT [14] and HVO (Mar Martinez-Pastor, unpublished data). The Ribo-Zero kit used biotinylated RNA probes designed to deplete abundant rRNA transcripts from bacterial total RNA with streptavidin beads. We observed 100% removal of rRNA from HVO total RNA samples (**Fig. 2**). In contrast, removal from HBT was variable, with a median rRNA value of 35% (range 18.7% -46.4%; **Fig. 2;** Table S1), at a level which allowed analysis of differential expression [14]. Because RNA-seq transcriptomic profiling studies across halophilic archaea are valuable to understand responses to environmental perturbation, we were hence motivated to find a suitable replacement capable of matching or bettering this performance across four model species of halophiles routinely used in our lab (HBT, HVO, HFX and HAH, abbreviations listed in **Table 1**).

**Fig 2.**
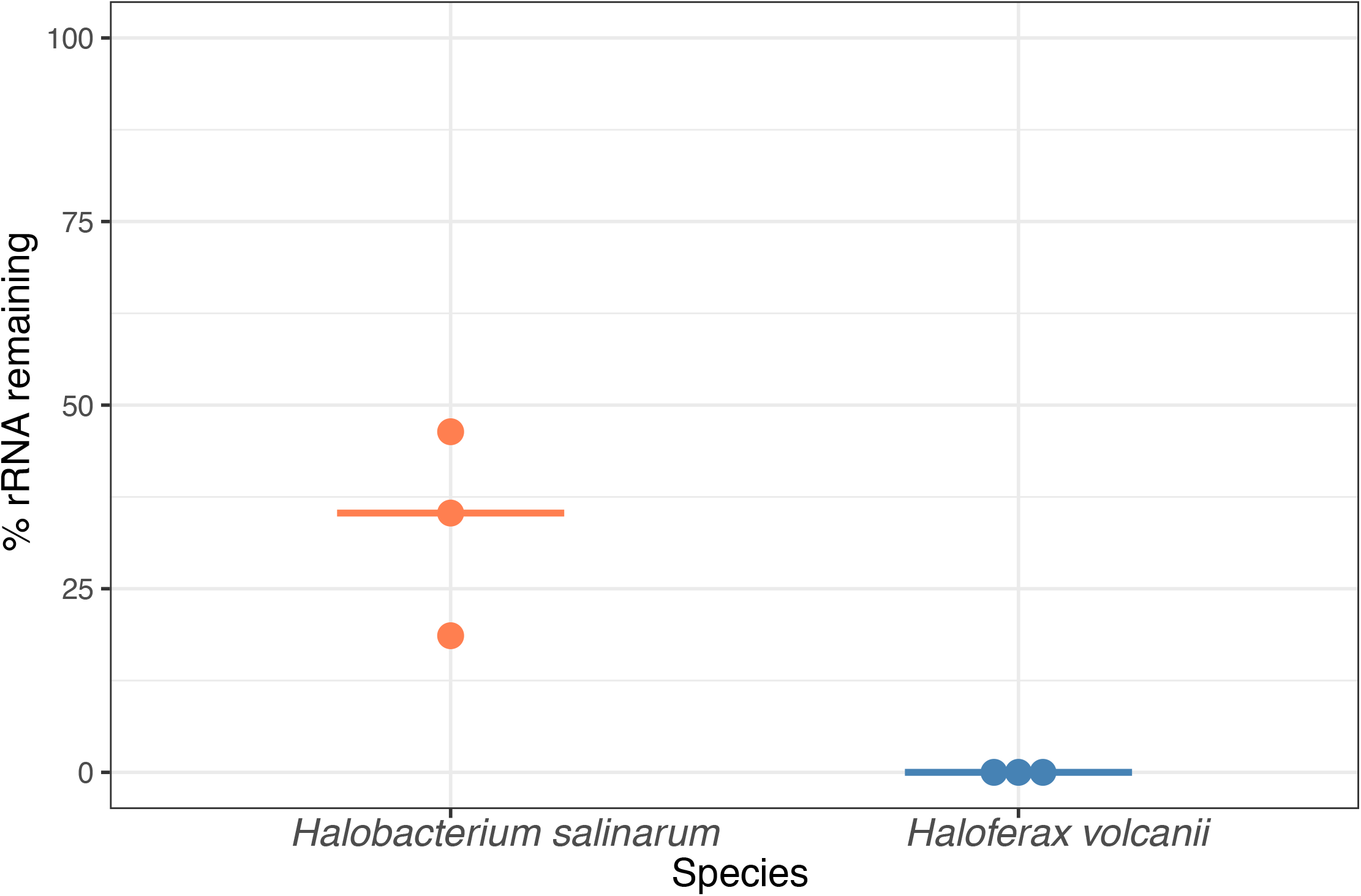
Percentage of rRNA remaining in halophile RNA by using the discontinued Ribozero kit (RZ). Each dot denotes one sample, with light orange dots representing *Hbt. salinarum* and blue dots representing *Hfx. volcanii* samples. Horizontal bars represent the median % rRNA remaining.

### Testing new rRNA depletion strategies on total RNA samples from *Halobacterium salinarum* (HBT)

We first began with a quantification of rRNA removal in HBT to allow continuation of ongoing differential gene expression experiments [34]. We used three enzymatic digestion-based rRNA depletion approaches from the following commercial kits (details in **Table S1** and Methods): (a) NEBNext Bacterial rRNA Removal Kit (probes included, abbreviated throughout as “NEB-B”); (b) NEBNext rRNA Core Depletion Reagent Set (with user-designed probes specific for HVO, method abbreviated throughout as “NEB-HVO”); and (c) the newly released Ribo-Zero Plus kit from Illumina (includes probes allowing universal depletion across bacteria and eukaryotes, “RZ+”). Following rRNA removal, resultant RNA samples were subjected to Next Generation sequencing, and the number of rRNA reads removed was quantified as compared to an untreated RNA control (Methods).

We observed that ∼95% of reads from sequenced untreated RNA correspond to rRNA (**Fig. 3**, Table S1). RZ+ treatment achieved a negligible reduction of rRNA to ∼92%. A slightly more substantial reduction was seen with the NEB-B method, with a median remaining rRNA percentage of 86%. Of these methods, the best results were obtained using NEBNext with customized probes designed to bind HVO rRNA sequences (NEB-HVO), although high levels of rRNA still remained (median remaining rRNA 80.5%, range 63% to 86%). We note that using no removal, RZ+, and NEB-HVO methods result in a range of ∼1.5-3.6M reads mapping to non-rRNA genes per sample (with 12 total samples run on one lane, **Table S1**). Based on our power analysis using online tools [38], this level of sequencing depth would require 5-6 biological replicates for reliable detection of 75% of differentially expressed genes (FDR < 0.05, log fold change >= 2.0) (**Fig. S1**). Since this depth was achieved with 12 samples multiplexed per lane, a requirement of 4-5 samples of each type would restrict RNA-Seq experimental design to a single comparison (for example, two genotypes in one condition or two conditions for the same genotype) per lane. Hence, the inefficient rRNA removal severely limits the extent to which samples can be multiplexed, increasing costs even in modern high-throughput sequencing instruments used here (**Table S1**).

**Fig 3.**
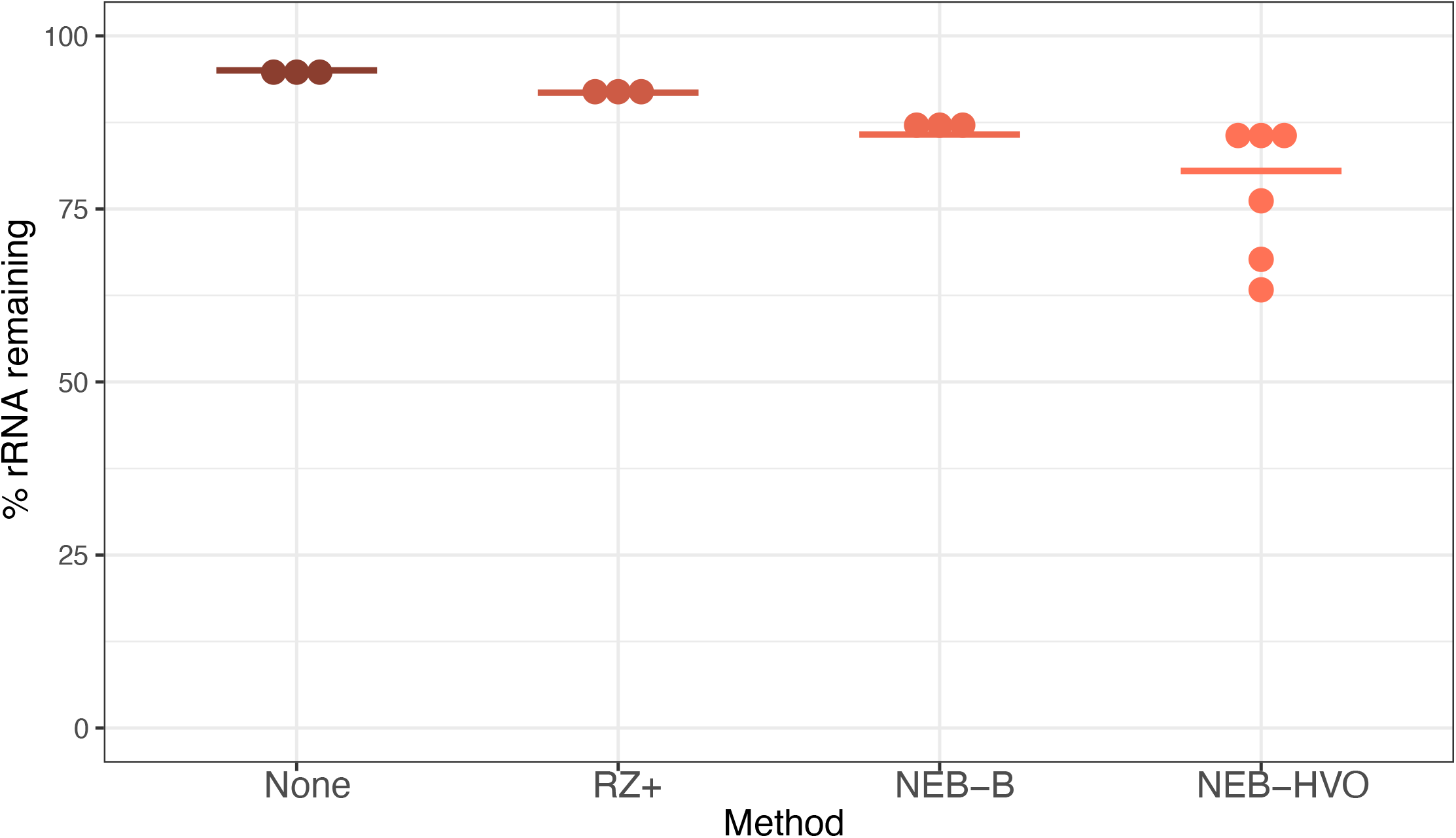
rRNA removal using alternative methods in *Hbt salinarum*. Each dot represents percentage of counts mapping to rRNA genes after using no removal (brown), New Ribozero kit (RZ+, dark orange), NEBNext kit with bacterial probes (NEB-B, orange), and NEBNext kit with HVO probes (NEB-HVO, peach). Horizontal bars represent the median value.

We hypothesized that poor rRNA removal may stem from either the incomplete RNase H digestion or the imperfect sequence match between the HVO rRNA probes used in the NEB-HVO method and the rRNA genes of HBT. To test the efficacy of RNAse H digestion, we carried out this digestion over 30 minutes (manufacturer protocol) and 120 mins (extended digestion) using the NEB-HVO method. Each digestion time used the same extracted RNA sample (split into two different aliquots for digestion), and was performed in biological triplicate within the same sequencing batch. A marked improvement in rRNA removal is seen in the 120-minute digestion (**Fig. 4A**), with 75% median rRNA remaining, as compared to 85% for the 30 minute samples. However, when comparing the results between different batches of sequencing, we found that the batch effect was stronger than the RNAse effect: 30 minute RNAse H digestion from a different batch produced a rRNA range of 63-76% (median 68%), better than even the 120 minute digestion from the first batch. Hence, while longer RNAse H digestion could potentially improve rRNA removal, this effect is inconsistent.

**Fig 4.**
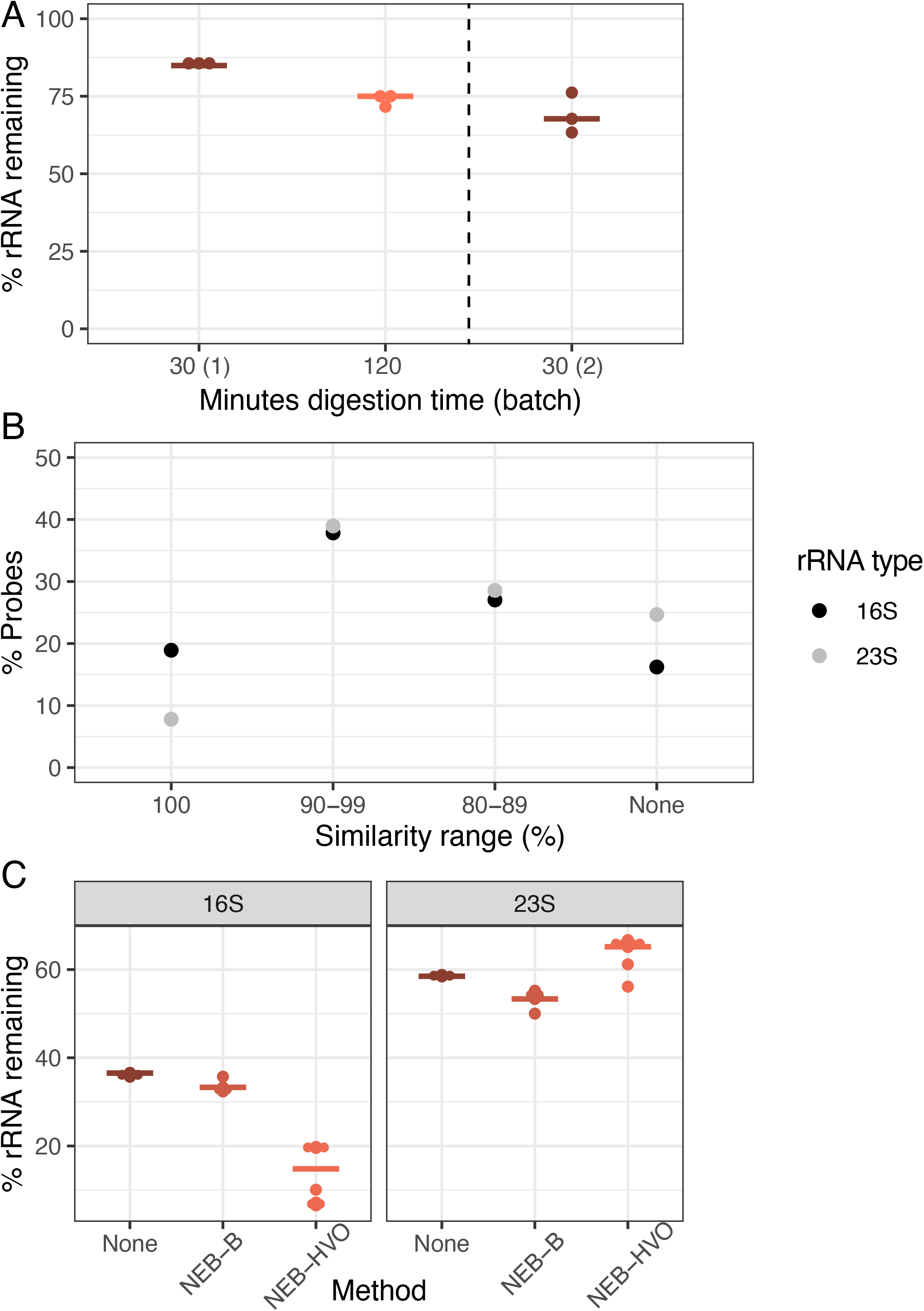
Increasing RNAse digestion time is less important than probe sequence identity for efficient rRNA removal. (**A**) Dotplot showing percentage of counts mapping to rRNA genes after using NEB-HVO method on HBT total RNA samples after 30 minutes (brown) or minutes 120 (light orange) of RNAseH digestion. “NEB30 (2)” samples to the right of the dotted line were processed and sequenced in a different batch. Horizontal bars represent the median value. (**B**) Percentage of custom-designed HVO probes classified into 16S (black) and 23S (grey). Levels of sequence identity of HVO probes with *Hbt. salinarum* (HBT)16S and 23S rRNA genes are shown on the X-axis, whereas percentage of total probes at each sequence identity level is shown on the Y-axis. (**C**) Percentage of total reads mapping to either 16S (left panel) or 23S rRNA (right panel) genes of HBT using 3 different rRNA removal methods - none (brown), NEB-B (dark orange), NEB-HVO (light orange).

Based on these results, we then tested the hypothesis that this relatively poor rRNA removal (compared to the discontinued RZ method) was associated with sequence mismatches between probes and rRNA. Using the NEB-HVO method, we observed that the probe sequences custom-designed for HVO rRNA matched HBT 16S rRNA sequences better than to 23S probe sequences (**Fig. 4B, Table S3**). 19% of 16S HVO probe sequences had 100% identity with HBT 16S rRNA, compared to only 8% for 23S rRNA. Conversely, 25% of 23S probe sequences shared no sequence similarity with HBT 23S rRNA, while this was only 16% for 16S. Corresponding with these different levels of sequence identity, we observed that 16S rRNA removal was more effective (∼15-35% remaining) than 23S rRNA removal (∼55-65% remaining, **Fig. 4C, Table S3**). Bacterial probes (included in the NEB rRNA removal kit) have very different rRNA sequences from those of archaea. Thus, as expected, the NEB-B method showed 16S and 23S rRNA remaining comparable to the no-removal control (**Fig. 4C**). Hence, there is a strong relation between probe sequence and rRNA removal, with even slight increases in probe specificity (**Fig. 4B**), resulting in profound differences in rRNA removal (**Fig. 4C**).

We conclude from these experiments that the NEBNext Core Reagent Set kit with probes custom-designed for the related species HVO (NEB-HVO) is the best of the reagent kits that we tested for HBT rRNA removal. RiboZero Plus (RZ+) and NEBNext Bacterial kit using the bacterial probes (NEB-B) led to less efficient rRNA removal for HBT. We expect that targeting custom probes specifically for HBT would likely result in better rRNA removal.

### Species-specific probe methods efficiently remove *Haloferax volcanii* (HVO) rRNA

Having shown the importance of probe sequence specificity, we next tested two different methods with rRNA probes targeted to HVO against HVO total RNA samples: (a) NEBNext Core Reagent Set (“NEB-HVO” method); and (b) the siTools RiboPool kit (“rP-HVO”). Unlike the enzymatic NEB-HVO method, rP-HVO uses streptavidin-based removal of rRNA hybridized to biotinylated probes. For both methods, we used probes custom-designed to be specific to HVO rRNA sequences (see Methods). We observed that both methods achieved nearly complete rRNA digestion: median values of 0.008% and 0.000008% rRNA remaining were observed using NEB-HVO and rP-HVO methods, respectively (**Fig. 5**; **Table S1**). These results with near-complete rRNA depletion in HVO with species-specific probes is in line with the observations above: the limiting factor with these probe-based methods is the identity of probe sequences with target rRNA sequences. Overall, we found that using probes targeted to HVO with either method resulted in efficient and near-complete removal of rRNA from HVO samples, with the riboPOOL method resulting in nearly undetectable rRNA.

**Fig 5:**
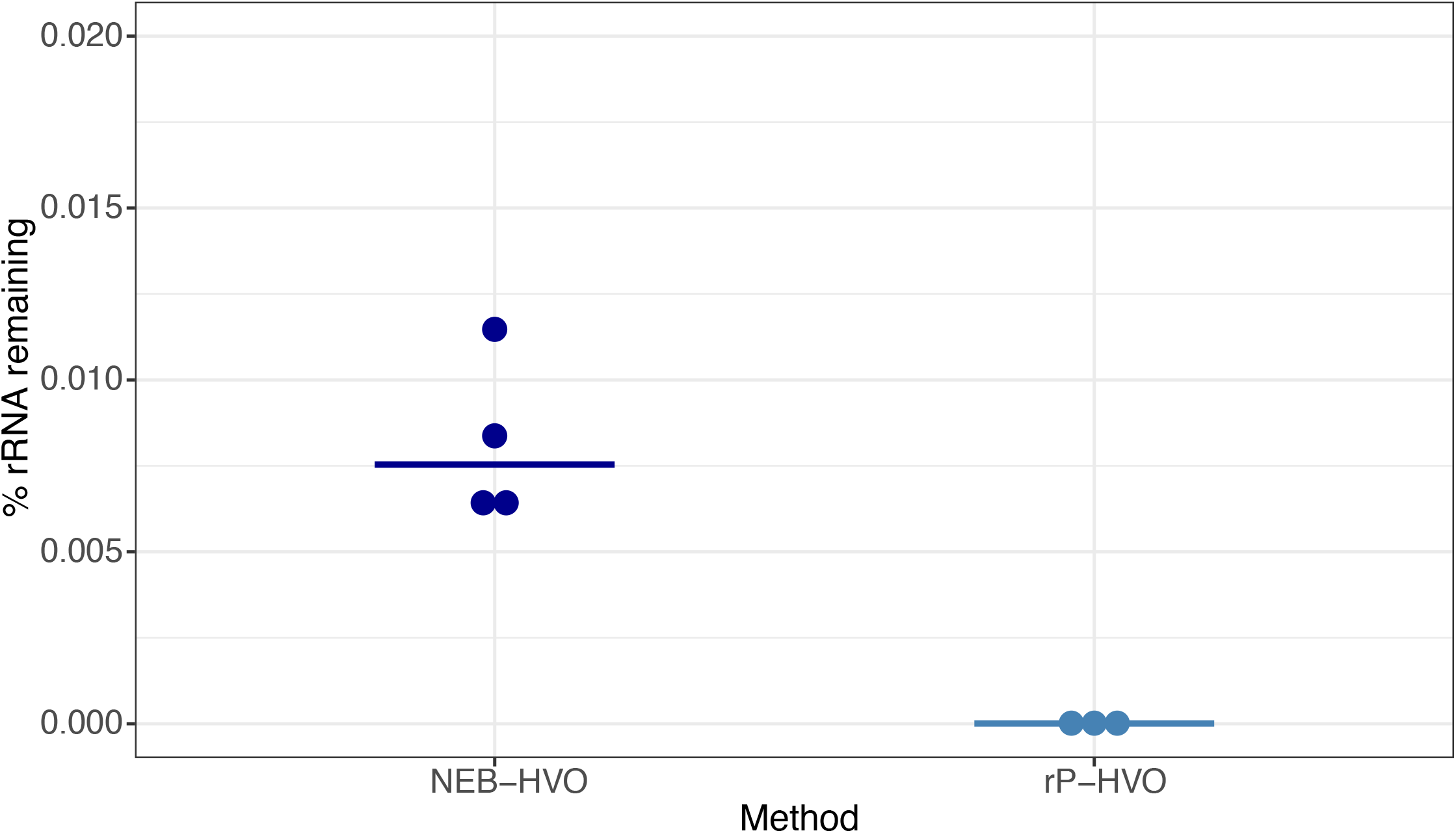
Species-specific probes efficiently remove rRNA from target species. Dotplots showing percentage of rRNA remaining after using probes with sequences specific for *Hfx. volcanii* (HVO) rRNA. Dark blue dots represent %rRNA remaining in individual replicate samples depleted with NEBNext Core Reagent Set (“NEB-HVO” method). Light blue dots represent %rRNA remaining in individual replicate samples depleted with the siTools RiboPool kit (“rP-HVO”). Horizontal bars represent the median value.

### siTools Panarchaea kit efficiently removes rRNA from diverse halophilic archaeal species

To expand our analysis to other model species, we then tested the siTools riboPOOL Panarchaea kit (rP-PA, **Table S1**, methods). The probe set associated with this kit is composed of high complexity pools of biotinylated DNA probes with sequences designed to deplete rRNA from a broad spectrum of archaea, including several classes of Euryarchaeota and Proteoarchaeota (https://sitoolsbiotech.com/ribopools.php). The Panarchaea riboPOOL probes have been shown to remove 99% of rRNA from *Sulfolobus solfataricus* and *Sulfolobus acidocaldarius* (https://sitoolsbiotech.com/pdf/microbes-ribopools-072021.pdf), but to our knowledge have not been published for euryarchaeal species like the four model halophiles of interest here. After using this kit for ribodepletion, we observed that all tested RNA samples across the four species contained <10% rRNA, with median values of 3.3%, 0.0002%, 0.04%, and 0.5% for HBT, HVO, HFX, and HAH, respectively (**Fig. 6, Table S1**). This extensive rRNA removal is more effective for HBT than for any previously tested methods (**Fig. 2, 3**), and equally as effective as NEB-HVO and rP-HVO methods for HVO (**Fig. 4, 5**). The other two species had not been previously tested, and no other RNA-seq results (other than for HFX small RNAs, which does not require rRNA removal [39]) are available for comparison in the literature. Taken together, these results demonstrate that the Panarchaea method (rP-PA) efficiently removed rRNA for four different model species of halophilic archaea.

**Fig 6.**
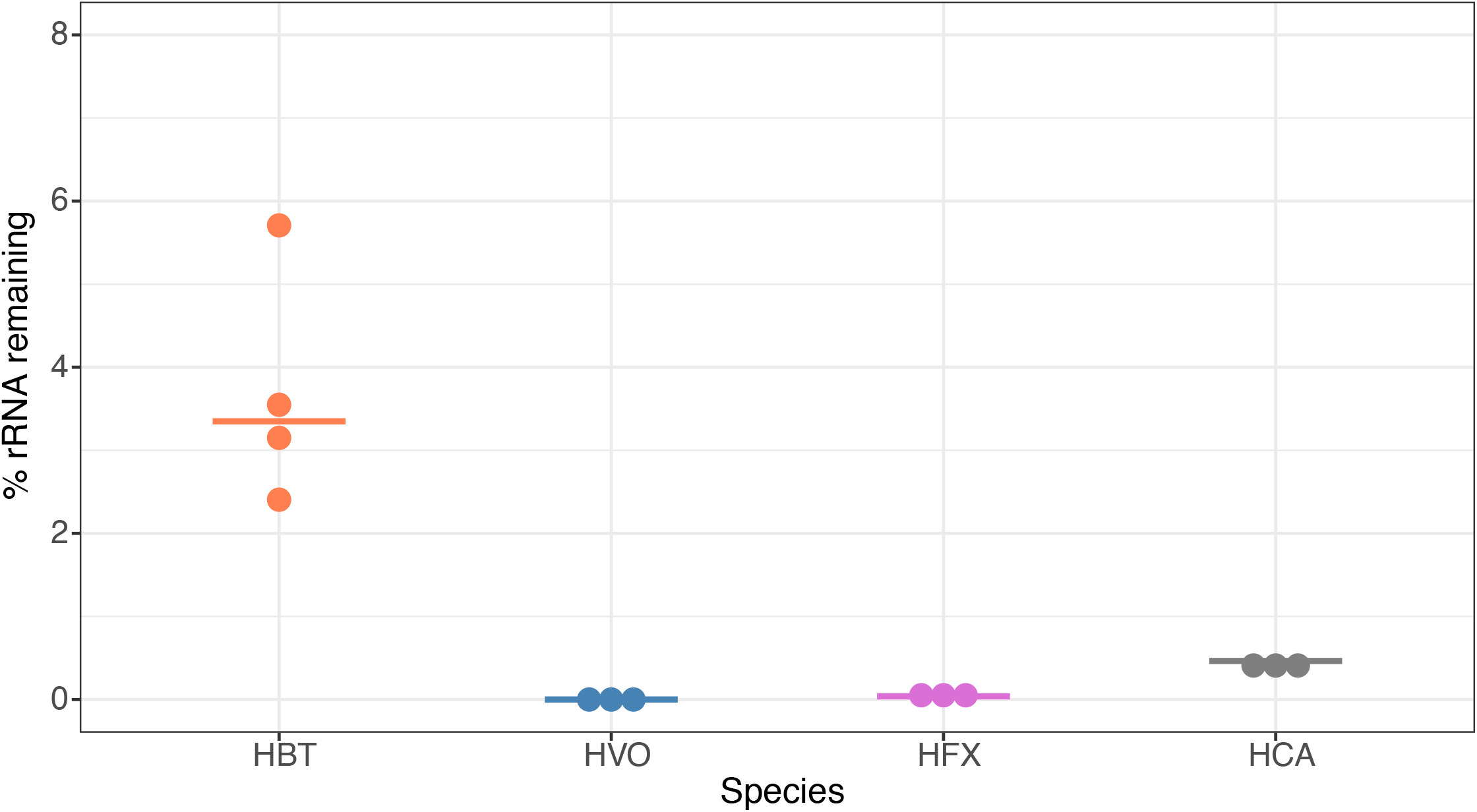
Panarchaea kit efficiently removes rRNA from total RNA across halophilic species. Dotplots showing percentage of remaining counts mapping to rRNA genes in *Hbt. salinarum* (HBT, orange), *Hfx. volcanii* (HVO, blue), *Hfx. mediterranei* (HVO, purple), *Hca. hispanica* (HCA, grey). Horizontal bars represent the median value of three biological replicate samples.

### Choice of removal method does not affect per-gene read counts

It was observed previously that using different rRNA removal methods can affect relative read counts of some non-rRNA genes [40,41]. We therefore tested whether rRNA removal and the choice of removal method changes the relative levels of mRNA. We calculated gene counts from each sample as a percentage of the total (non-rRNA) counts from that sample, and correlated these relative counts obtained from different rRNA removal techniques (see Methods, **Fig. 7**). We observed strong correlations of normalized relative counts of non-rRNA genes among different rRNA removal methods, as well as with untreated total RNA data for HBT (**Fig. 7A**). The Pearson’s correlation coefficients between per-gene normalized read counts across different methods used on HBT were in the range 0.91-0.99, with an average value of 0.95. Correlation with control (untreated) samples was >0.92. Similar results were seen with HVO: 0.94-0.99, average 0.97 (**Fig. 7B, Table S4**). Based on this analysis, we conclude that rRNA removal and the choice of removal method does not change the number of reads on a per-gene basis in halophilic archaea. These rRNA removal methods can therefore be used for downstream applications such as differential gene expression analysis.

**Fig 7:**
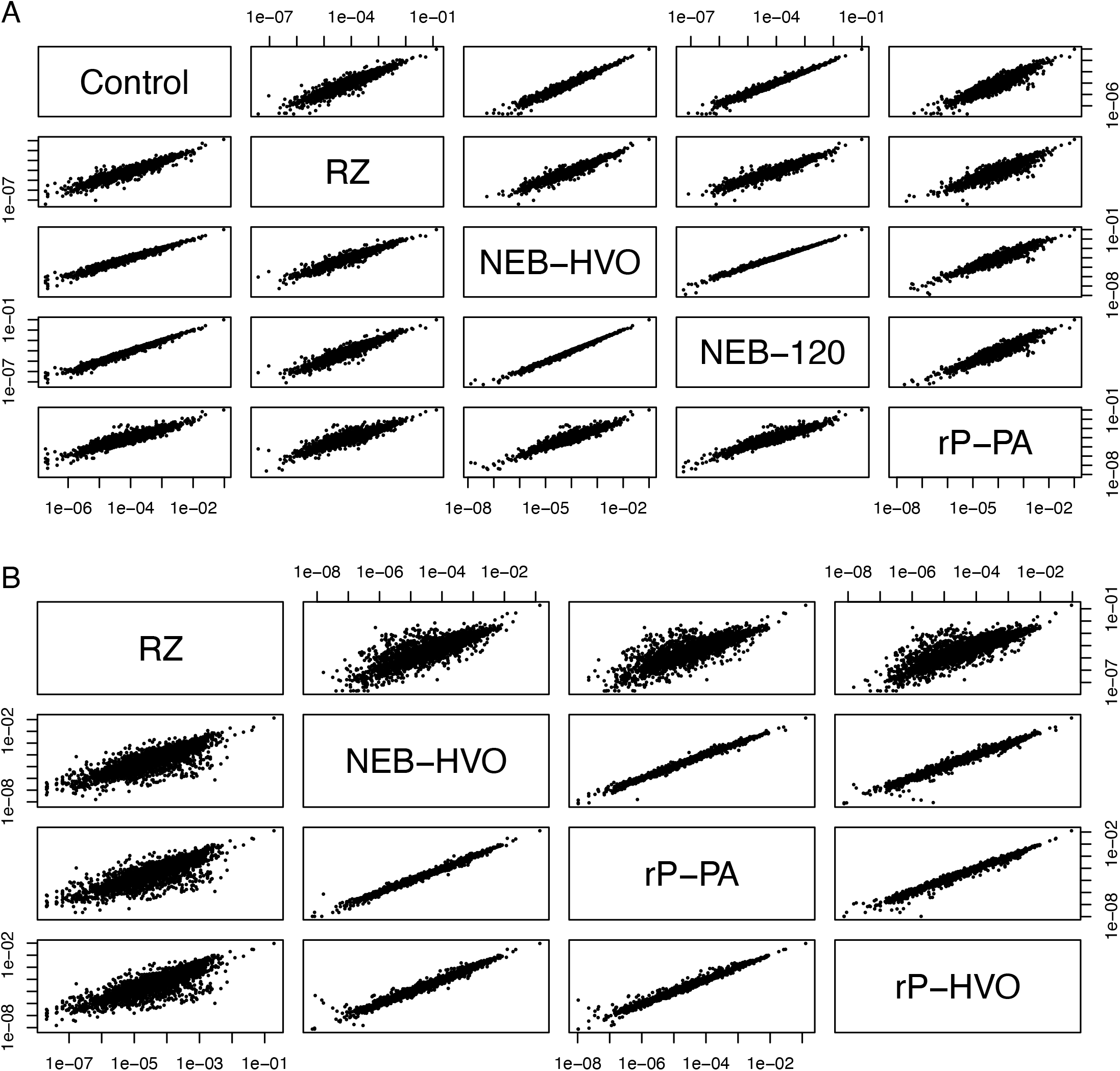
Choice of removal method does not affect relative abundance of mRNAs. Correlations between relative abundance of each gene after different rRNA removal methods in (**A**) *Hbt. salinarum* (HBT) and (**B**) *Hfx. volcanii* (HVO). Each dot represents the percent of total normalized reads for each gene (see Methods section). Methods shown here are “Control” (no removal), “RZ” (using discontinued RiboZero kit), “NEB-HVO” (using NEBNext kit with custom HVO probes), “NEB-120” (NEBNext kit with custom HVO probes and 120 mins of RNAse digestion), “rP-PA” (siTools riboPOOL method using Panarchaeal probes), and “rP-HVO (siTools riboPOOL method using HVO-specific probes).

### Utility of rRNA removal is seen in counts of non-rRNA genes

Previous studies have suggested a minimum sequencing depth of two [19] to ten [20] million reads per sample for obtaining reproducible results for differential expression. On a per-gene basis, 5 reads is considered a threshold below which differential expression analysis is unreliable [18]. We sought to understand how rRNA removal affects transcript detection using data from HBT, from which we have data for a wide array of rRNA removal methods (including no removal), and a large range of rRNA remaining in sequenced samples (2%-95%). Across these samples, we calculated the number of annotated genes with no mapped reads as well as <5 mapped reads. For consistency, we only considered samples that had been sequenced on the same machine (NovaSeq6000). We observed that more complete rRNA removal generally leads to increased detection of genes (**Fig. 8)**. All numbers that follow are median values, obtained from **Table S1**. For untreated RNA (∼95% rRNA), ∼320 genes showed no reads, and this reduced to ∼312 for RNA treated with NEB-B (∼86% rRNA) and further to ∼307 with NEB-HVO (∼68% rRNA). The Panarchaea kit (rP-PA) reduced the number of undetected genes to ∼298 (∼3% rRNA). For genes with < 5 counts detected, there was a more dramatic change, from ∼385 genes for untreated RNA, but only ∼310 for Panarchaea kit. This trend held true even though the total number of reads for all genes (including rRNA) had relatively similar median values of ∼30M and ∼27M reads, respectively (**Table S1**), suggesting that the improved detection of lowly expressed genes is associated with more complete rRNA removal rather than deeper overall sequencing of the samples. These above results indicate that better rRNA removal improves per-gene counts and detection of lowly expressed genes, which is important for making accurate statistical conclusions regarding differential expression. rRNA removal enables increased multiplexing and therefore reduced cost for RNA-seq experiments.

**Fig 8.**
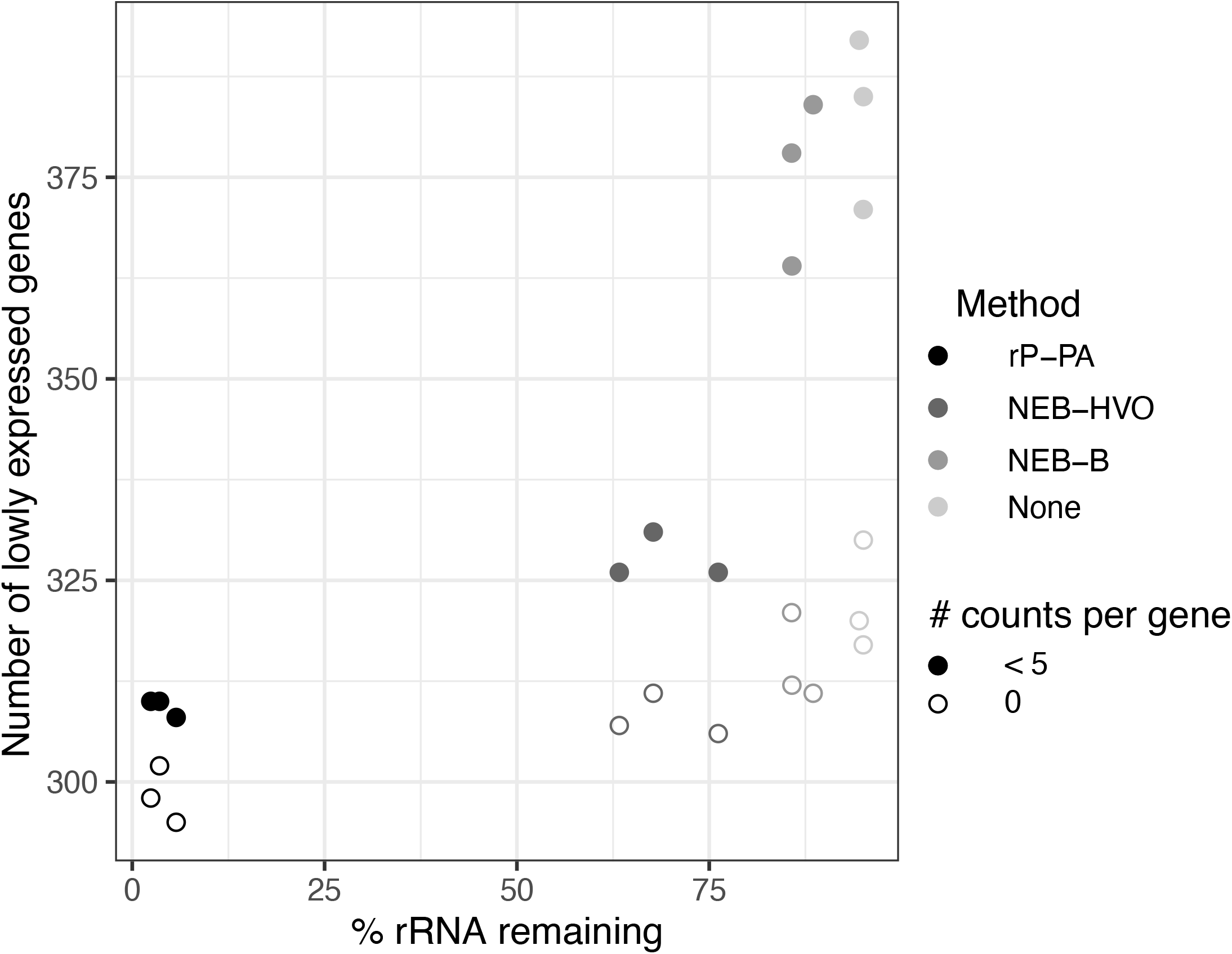
More complete rRNA removal leads to increased detection of lowly expressed genes. Number of genes with zero (open circle) and <5 (filled circle) reads detected in sequencing samples treated with different rRNA removal methods in *Hbt. salinarum*. The darker the circle color, the more complete the rRNA removal for each method: riboPOOL Panarchaea (rP-PA, black); NEBNExt with HVO-specific probes (NEB-HVO, dark grey); NEBNext with bacterial probes (NEB-B, grey); no removal (none, light grey).

## CONCLUSIONS AND DISCUSSION

The main technical challenge for prokaryotic transcriptomics is the low ratio of mRNA:rRNA. Historically, different methods have been used to eliminate rRNA without biasing mRNA reads: from digestion with exonucleases that preferentially degrade rRNA relying on 5’ monophosphate; to subtractive hybridization that captures rRNA binding to antisense oligonucleotides [42,43]; to poly(A) tail addition to discriminate rRNA or reverse transcription with rRNA primers followed by RNaseH digestion [30,44]. However, none of these methods has been successfully utilized for haloarchaea. Until the end of 2018, the Ribo-Zero kit from Illumina, based on sequence-specific biotinylated probes that hybridize with a pool of microbial rRNA sequences and then selectively remove the hybrids using streptavidin-coated magnetic beads, enabled removal of ∼70% of rRNA for several archaeal species [13-17]. After this commercially available kit was discontinued, archaeal transcriptomics was at an impasse. Here we invested time to troubleshoot this problem, test, and directly compare newly available tools to help the archaeal community to move on with transcriptomic studies. Our investigation provides a guide for choosing a suitable application depending on the model organism or the combination of archaeal species of interest (e.g. communities, labs using multiple cultured species, metatranscriptomics).

We found that both RNAseH-based and biotin-based methods are efficient for rRNA removal. Certain commercially available kits from NEB and siTOOLs are most effective when probes are designed that target archaeal species of interest. For HVO, the RiboPool kit as well as the NEBNext kit with custom-designed probes that target HVO (**Fig. 5**) resulted in nearly complete rRNA removal. A similar number of total reads was observed after sequencing with no detectable bias in lowly expressed transcripts (**Fig 8**). In general, when using targeted kits, we found that the most important factor in determining rRNA removal efficiency was percentage identity of the target rRNA with the probe sequence (**Fig. 4, 5**). We found that the Panarchaea kit from siTOOLs provides very good rRNA depletion across all four species tested here (**Fig. 6**) and we anticipate that these can be effectively used for metatranscriptomics of archaeal communities. Targeted methods from both NEB and SiTOOLs as well as the Panarchaea probe set provide comparable performance to the discontinued RiboZero kit for HVO, with remaining rRNA close to 0. We further note that the Panarchaea kit exceeds the performance of the previous RZ method for HBT (**Table S1, Fig. 2 vs Fig. 6**). In the future, continuing to deposit raw RNAseq data from the archaeal community into online data repositories such as NCBI Gene Expression Omnibus is critical for progress in the area of transcriptomics, which would facilitate future efforts to predict rRNA removal success depending on probe sequence identity.

One of the most important advantages of choosing and efficient rRNA removal method is analyzing differential expression of low-count genes. In the current study, we show that efficient rRNA depletion enables increased detection of lowly expressed genes (**Fig. 8**). This improvement in coverage of low-count genes enables correct statistical analysis of differential expression [19-21]. Accurate detection of lowly expressed transcripts is also important when using RNA-Seq to map the transcriptome [12,45], including in metatranscriptomic protocols [44]. The rapid pace of discovery of new archaeal species [6,46] as well as the use of novel archaeal model organisms in lab will bring further challenges for transcriptomics experiments. However, the methods tested here provide sufficient flexibility to solve such challenges. For example, it is possible that newly identified archaeal species may encode rRNA sequences divergent from commercially available primer sets such as siTOOLs Panarchaea. The NEBNext Core Reagent Set using the custom probe design tool (https://depletion-design.neb.com/) would therefore be an appropriate choice in this case. Removal of rRNA enables increased detection of rare transcripts and extensive multiplexing. The methods tested here will therefore facilitate rapid progress in understanding the transcriptional response of a wide diversity of archaea to their environment.

## AUTHOR CONTRIBUTIONS

Conceptualization, MMP, SS, DNR, and AKS; methodology, MMP, DNR, and SS; software, SS; validation, SS and AKS; formal analysis, SS; investigation, SS and MMP; resources, AKS and DNR; data curation, SS and AKS; writing—original draft preparation, MMP and SS; writing— review and editing, SS, AKS, DNR, and MMP; visualization, SS and AKS; supervision, AKS; project administration, AKS; funding acquisition, AKS. All authors have read and agreed to the published version of the manuscript.

## DATA AVAILABILITY STATEMENT

Code used to analyze data associated with this study is freely accessible via https://github.com/amyschmid/rRNA_analysis. RNA-seq data is available through the National Center for Biotechnology Information Gene Expression Omnibus (NCBI GEO) database at accession GSE200776.

## ACKNOWLEDGEMENTS

We thank Schmid lab members for the comments on the data analysis and manuscript text, especially Rylee Hackley and Alex Phillips. We thank members of the haloarchaeal research community for ongoing discussions regarding the challenges with rRNA removal, especially Jocelyne DiRuggiero, Ron Peck, and Alexandre Bisson.

## FIGURE LEGENDS

**Figure S1** - Analysis of sequencing depth, number of biological replicates, and detection of differentially expressed genes (2-fold differential expression) using the online tool Scotty. Squares with red dots are predicted to have <75% detection of differentially expressed genes.

