## Supplementary figures and images for "Comparative Analysis of rRNA removal methods for RNA-seq Differential Expression in Halophilic Archaea"

### Figure S1

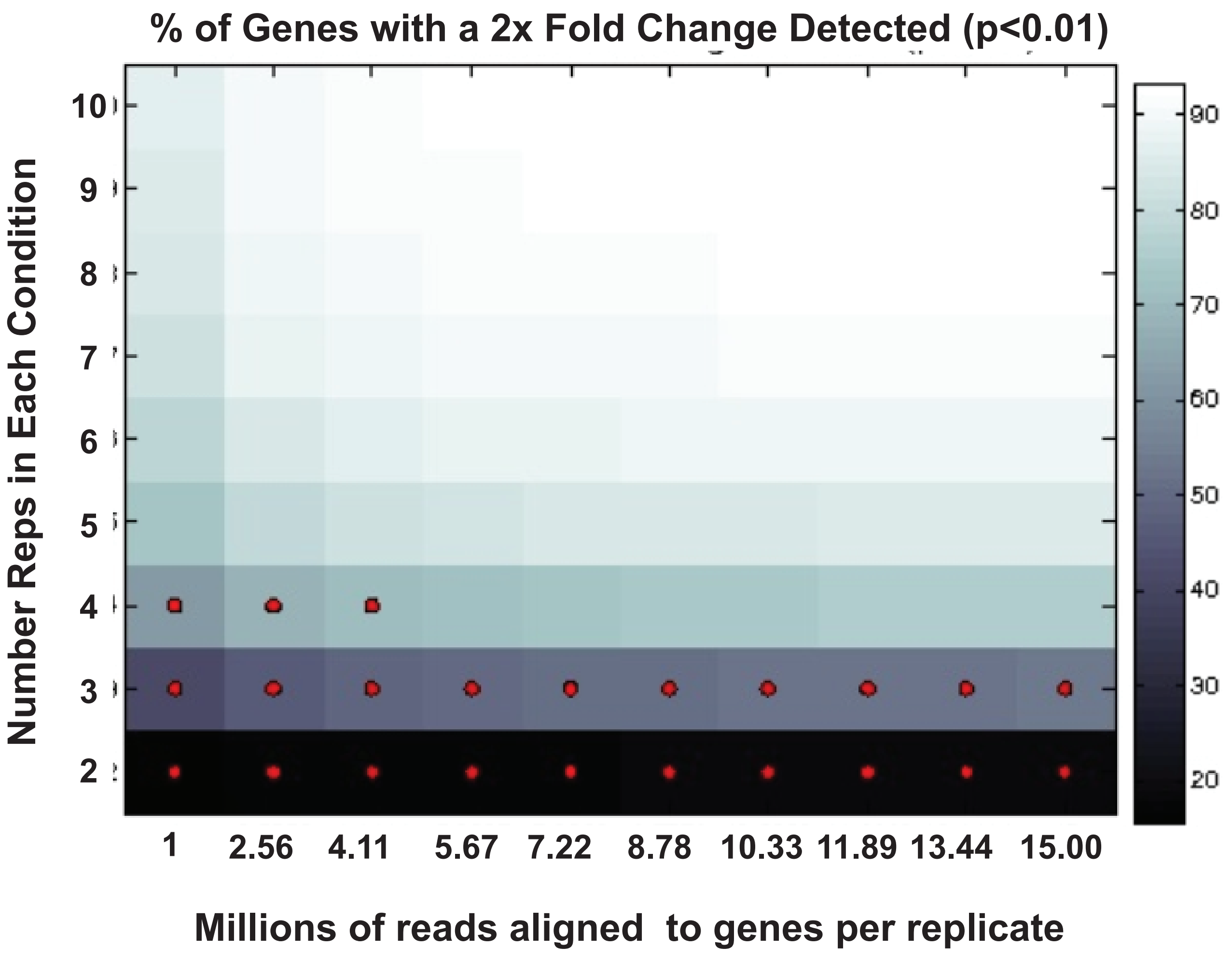
